# Elavl1 is dispensable for appendicular skeletal development

**DOI:** 10.1101/2024.09.23.614008

**Authors:** Rohini Parsha, Satya K. Kota

**Affiliations:** Department of Oral Medicine, Infection and Immunity, Harvard School of Dental Medicine, Harvard University, Boston, USA

## Abstract

Elavl1/HuR is a RNA binding protein implicated in multiple developmental processes with pleiotropic roles in RNA life cycle. Loss of Elavl1 is incompatible with life with early embryonic loss of Elavl1 in epiblast cells being lethal with defects in placental branching and embryonic tissue growth. Postnatal global deletion of Elavl1/HuR results in lethality with atrophy in multiple tissues mainly due to loss of progenitor cells. However, roles of Elavl1 specifically during embryonic limb development is not well understood. Here we report that deletion of Elavl1 in limb bud mesenchyme in mouse did not reveal any abnormalities during embryonic development with normal development in pre- and postnatal limb skeleton. Analyses of skeletal patterning, morphogenesis and skeletal maturation including skeletal elements in stylopod, zeugopod and autopod during development did not reveal any significant differences between long bones from control and Elavl1 conditional knockout animals. Our study indicates differential dependency and susceptibility to loss of Elavl1 in different stem cell lineages with its functions being dispensable during limb development.

## Introduction

RNA binding proteins play very important roles in maintaining organismal homeostasis as well as during disease resolution^1^. Elavl1 (embryonic lethal-abnormal vision like protein 1) / HuR (Human antigen R) protein has important roles in controlling the cellular levels and stability of multiple AU or U rich RNA transcripts^2^. Elavl1 regulation of RNA stability generally occurs via interactions with AU rich elements (AREs) in the introns and / or 3′ untranslated regions (UTRs) of messenger RNAs^3,4^. Interactions between ARE containing mRNAs in the nucleus and Elavl1 in general lead to stability by preventing access of these mRNAs to RNA binding proteins that recruit additional proteins involved in RNA decay pathway^5 6 7^. In humans and other mammals, Elavl family comprises of four members including ubiquitously expressed HuR/Elavl1 and developmentally regulated HuB/Elavl2, HuC/Elavl3 and HuD/Elavl4^8-10^. All of the Elavl family members contain RNA recognition motifs (RRMs) that bind to U and AU rich sequences (AREs) in RNAs predominantly in the UTR regions^11,12^. Ubiquitous distribution of Elavl1 points to fundamental roles in posttranscriptional regulation of RNA in multiple tissues during tissue homeostasis and development.

Elavl1 and its homologues regulate RNA metabolism and stability in many eukaryotic species from Drosophila to Humans^13^. Deletion of HuR in Drosophila resulted in embryonic lethality with developmental defects including in nervous system^14^. In mice, both pre- and postnatal deletion of Elavl1 leads to lethality. Global and epiblast specific deletion of Elavl1 using Sox2-cre led to defective development with embryos showing defects in skeletal, splenic development and embryonic lethality^15^. Postnatal global deletion of Elavl1 using tamoxifen-inducible Cre mice at 8 weeks of age also resulted in atrophy of multiple tissues and lethality within 10 days of depletion of Elavl1. Further, apoptosis of progenitor cells residing in multiple tissues including bone marrow, thymus and intestine was detected upon depletion of Elavl1 postnatally^16^. Recently, Elavl1 was also found to be essential in developing cranial neural crest^17^ and as a key protein for control of hepatic metabolic homeostasis^18^. In vitro, in cultured Bone Marrow Stomal Cells (BMSCs), Elavl1 adversely affected osteogenic differentiation by controlling stability and cellular expression levels of several ARE containing mRNAs associated with ECM organization^19^. In contrast to the above known roles in multiple tissue specific functions, very little is known about the role of Elavl1 during skeletal development. With the aim of understanding the role of Elavl1 during limb skeletal development, we generated Elavl1 conditional knockout mice using Prx1 cre that is expressed in limb bud and a subset of cranial mesenchyme. Our results indicated that Elavl1 is expressed in developing limbs however, loss of its expression in a specific and conditional manner in Prx1-cre expressing cells was compatible with life and led to normal skeletal development as analyzed across multiple stages during and post embryonic development.

## Methods

### Mouse crosses

All experiments involving mice were approved by the institutional IACUC. *Prx1* cre (B6.Cg-Tg(Prrx1-cre)1Cjt/J, Strain #:005584) male mice that have transgene, Prrx1 promoter/enhancer sequence directing Cre recombinase expression in early limb bud mesenchyme and a subset of craniofacial mesenchyme and Elavl1 floxed^16^ (B6.129-Elavl1^tm1Thla^/J, Strain #:021431) mice with loxp sites flanking exons 2-5 were obtained from Jackson labs. Mice were housed in sterile cages with access to food and water ad libitum, 12 hours of light and 12 hours of dark cycle. Male Prx1 cre (prx1-cre) and female Elavl1 foxed mice were bred to generate Prx1-cre; Elavl1fl/fl mice. Prx1-cre, Elavl1fl/fl and Prx1-cre; Elavl1fl/+ mice were used as controls. Genotyping was performed with the DNA isolated from ear punch tissues using the primers, Elavl1, Fw-CTC TCC AGG CAG ATG AGC A and Rev-TAG GCT CTG GGA TGA AAC CT.

### Embryo collection and skeletal preparations

Time-mated pregnant mice were euthanized at indicated timepoints to collect the embryos at different developmental stages. Alcian blue and alizarin red staining of skeletal preparations were performed essentially as described^20^. Forelimb, hindlimb and cranial skeletons were dissected from the whole mount skeletal preps and imaged separately. Analysis was performed for the presence and maturation status of individual bones in stylopod, autopod and zeugopod of fore and hind limb skeletal elements.

### Realtime qRT-PCR

Total RNA was prepared from hindlimbs collected from 17.5 dpc embryos using RNeasy kit (Qiagen). DNA contamination was eliminated by DNase I digestion. Complementary DNA was prepared using multiscript reverse transcription system (Applied biosystems). The reverse transcribed cDNA was subjected to qPCR using a Sybr green based detection system (Qiagen). Relative levels of transcripts were normalized to NAT1 or beta-actin or 18rDNA levels and quantified based on 2^−ΔΔCT^ method. A minimum of 3 limbs from conditional knockout (cKO) or controls were analyzed. The primer sequences used will be available upon request.

### Analysis of Elavl gene expression and open chromatin at regulatory regions

Transcripts per million (posterior_mean_count) values for Elavl1, Elavl2, Elavl3 and Elavl4 were extracted from the poly A plus RNA sequencing data from Encode^21,22^ (ENCSR098WGB, ENCSR407MLM, ENCSR902MLV) that were obtained from 10.5 dpc, 11.5 dpc, 12.5 dpc, 13.5 dpc, 14.5 dpc and 15.5 dpc limb RNA pooled from embryonic fore and hind limb tissues. DNAse seq sequencing data peaks pertaining to Elavl1 promoter and genic regions for 10.5dpc, 11.5dpc and 14.5 dpc limbs was obtained from Encode (ENCSR466MZF).

### Limb, femur and tibial length measurement

Lengths of left and right fore- and hind limb from control and Elavl1 conditional knockouts from four-week-old mice were measured along the stylopod, zeugopod and autopod using digital vernier caliper. Tibia and femurs were collected from mice hind limbs and proximo-distal lengths were measured using vernier caliper.

## Results

To understand the biological role of Elavl1 during limb skeletal development, Elavl1 gene expression was analyzed during mouse limb development between 10.5 dpc and 15.5 dpc from polyA plus RNA sequencing data from ENCODE. Elavl1 mRNA expression was detected in all of the developmental stages analyzed, with highest expression seen at 10.5 dpc limb bud stage. Compared to 10.5 limb buds, gradual decrease in Elavl1 transcript levels was observed between embryonic stages 12.5, 15.5 dpc (Fig-1A). Presence of Elavl1 mRNA levels correlated with open chromatin at the Elavl1 regulatory regions during the developmental stages (Fig-1B).

**Figure 1.**
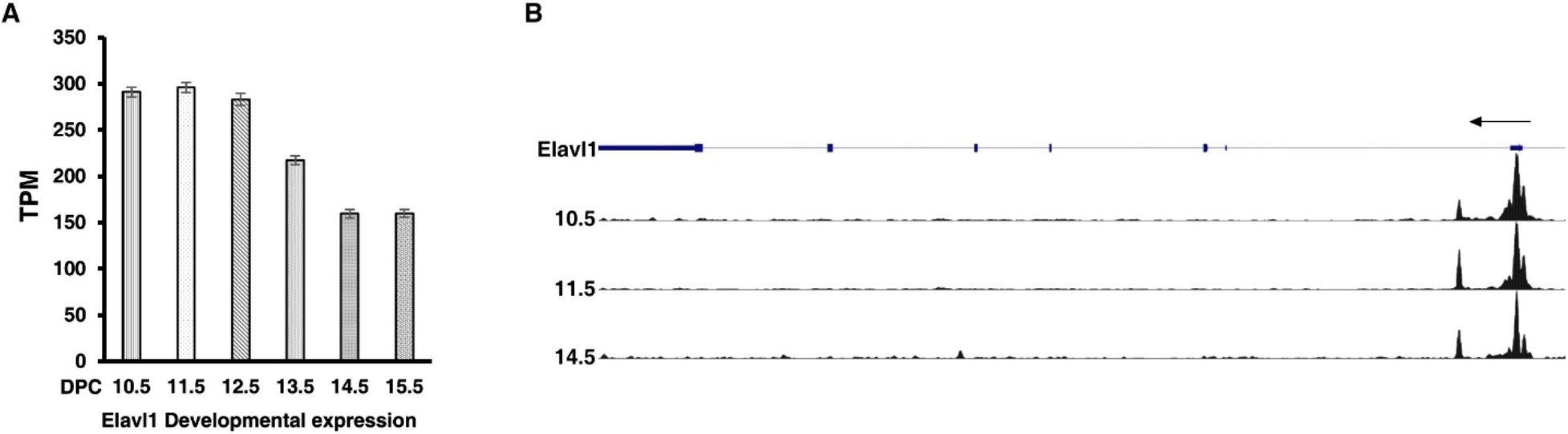
(A) RNA-seq analysis of messenger RNA levels of Elavl1 in limb RNA across developmental stages 10.5 dpc -15.5 dpc indicating expression seen across these developmental stages. (B) DNAse1-seq analysis of chromatin at the 10.5 dpc, 11.5 dpc and 14.5 dpc developmental stages shows DNAse I hypersensitive peaks indicating open chromatin near promoter regions of Elav1 gene.

As expression data clearly revealed the expression of Elavl1 in the developing limb bud, the gene was conditionally deleted from the early limb bud mesenchyme in Elavl1 floxed mice using cre recombinase expressed under the influence of Prx1 promoter/enhancer elements. Both control and Elavl1 conditional knockout (cKO) mice were obtained in expected mendelian ratios and survived to adulthood. Expression analysis of RNA from control and Elavl1 cKO embryonic hindlimbs by RT-qPCR indicated significant reduction in Elavl1 mRNA levels in Elavl1 cKO embryonic limbs compared to littermate controls.

Next, to further understand the effects of Elavl1 depletion on appendicular skeletal morphology, morphological analysis of control and Elavl1 cKO 13.5 dpc embryos (Fig-2A), P1 pups (Fig-2B) and 4 weeks old mice (Fig-2C) were performed. Gross morphological analysis did not reveal any significant differences between control and Elavl1 conditional knockout embryos or adult mice. Quantitation of forelimb and hindlimb lengths from four-week-old mice also did not reveal any significant differences between control and Elavl1 cKO mice (Fig-2D). Further, skeletal preparations were made from 15.5 dpc and 18.5 dpc embryos to understand the differences in endochondral ossification between control and Elavl1 cKO embryos during development. We did not observe any defects in appendicular skeletal elements and their stage specific ossification patterns in 15.5 dpc (Fig-3A) and 18.5 dpc (Fig-3B) skeletal preparations from Elavl1 embryos compared to control embryos. All the skeletal elements in stylopod, zeugopod, and autopod both in fore limbs (Fig-3C) and hind limbs (Fig-3D) from 18.5 dpc were analyzed. No aberrations in skeletal elements including scapula, humerus, radius and ulna of forelimb, femur, tibia, fibula were observed in Elavl1 conditional knockout limbs, clearly indicating the dispensability of Elavl1 during skeletal development.

**Figure 2.**
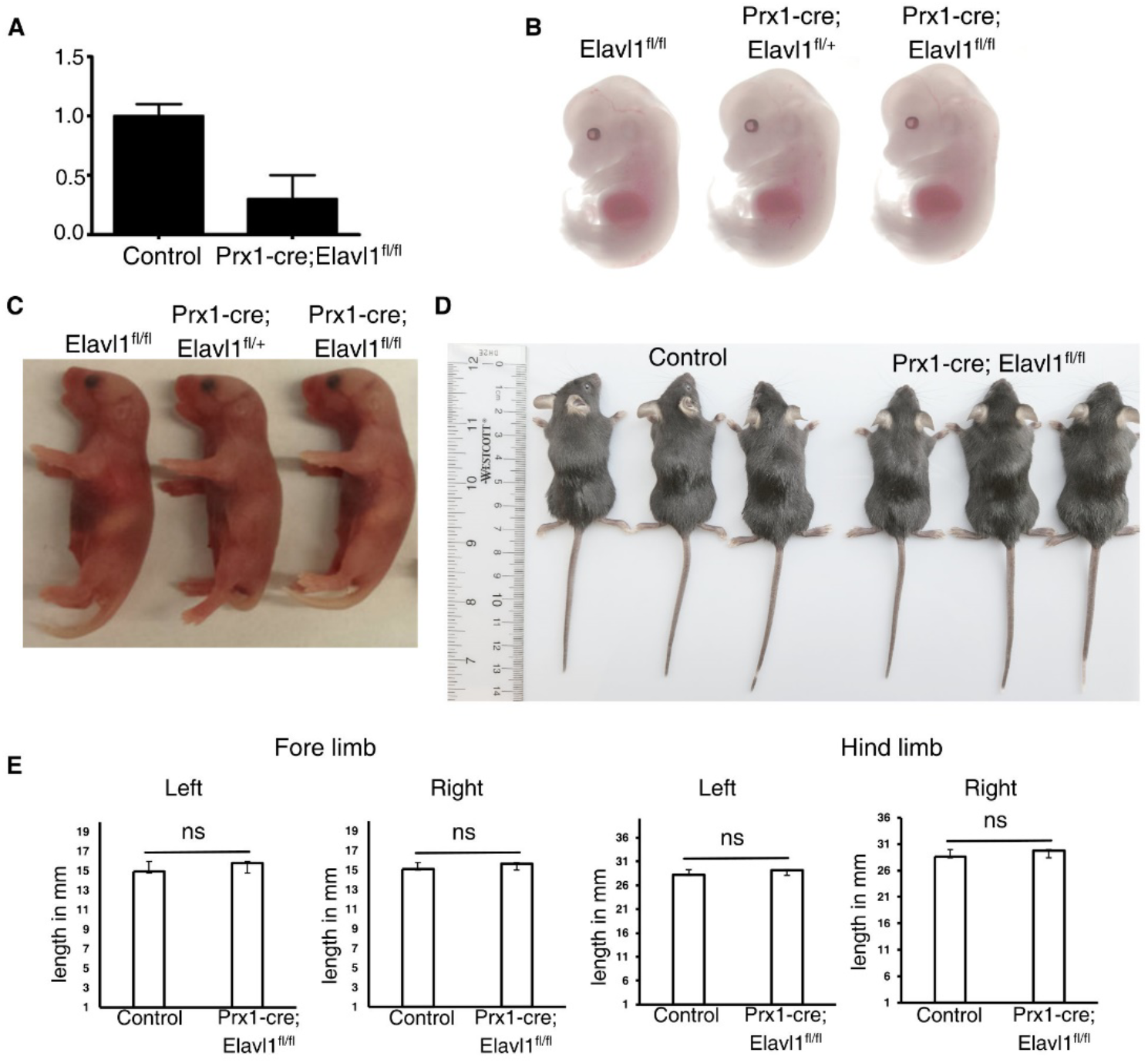
(A) RT-qPCR analysis of total RNA from control (N=3) and Elavl1 conditional knockout limbs (N=3) from 17.5 dpc embryos indicating significant downregulation of Elavl1 RNA levels in limbs upon Prx1-cre mediated deletion. Representative whole-body images of 13.5 dpc embryos (B), neonatal mice pups of P1 stages (C) of control (Elavl1^fl/fl^, Prx1-cre; Elavl1^fl/+^, N=8), Elavl1 conditional knockout (Prx1-cre; Elavl1^fl/fl^, N=7) showing no gross morphological differences during pre- and postnatal development. Whole body images of 4-week-old male control (N=3) and Elavl1 cKO (N=3) mice showing no gross morphological differences including limbs (D). Fore- and hind limb lengths in the proximal-distal axis in four-week-old male mice measured between control (n=3) and Prx1-cre; Ealvl1^fl/fl^ mice(n=3), ns: not significant

**Figure 3.**
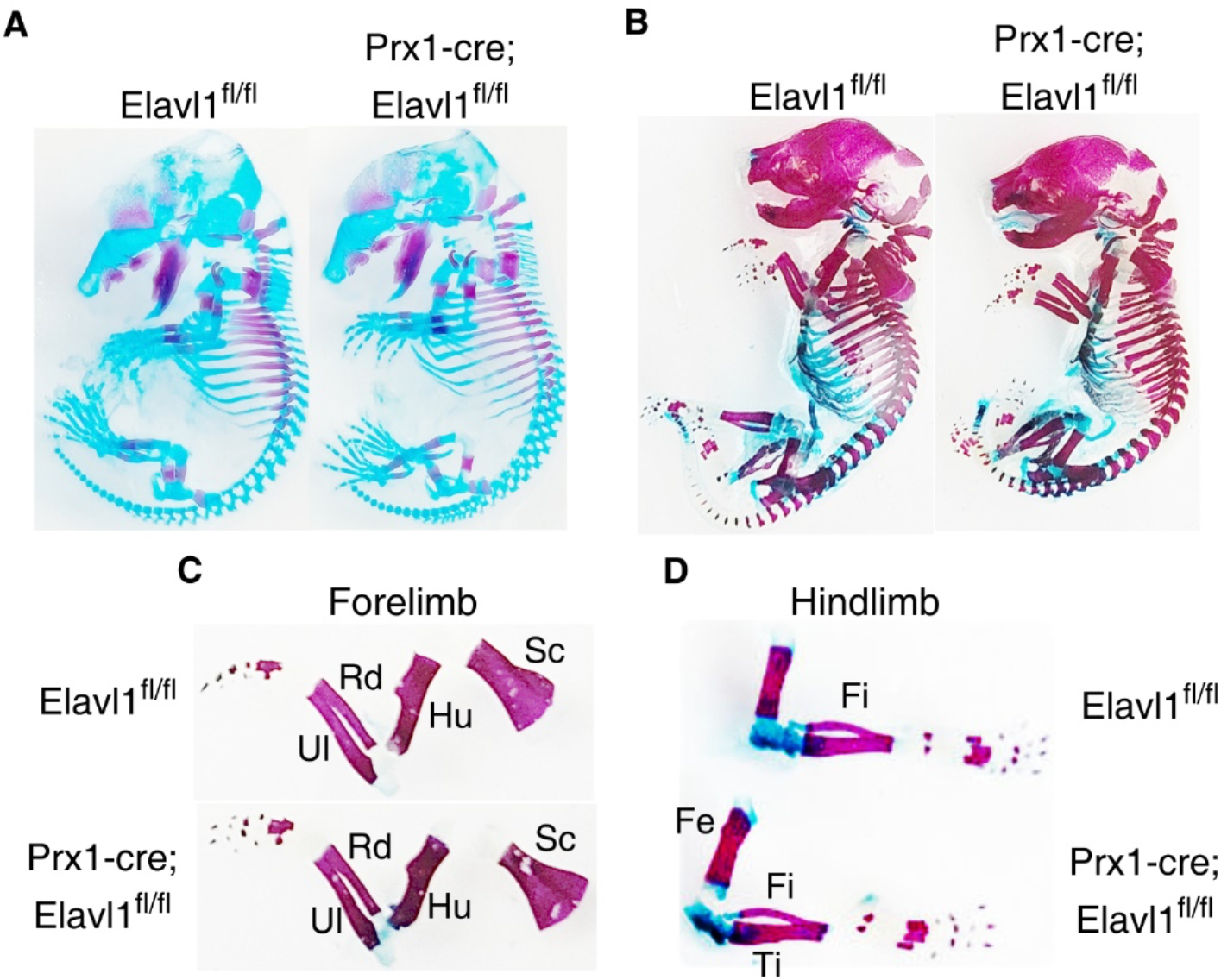
Representative whole body skeletal preps from 15.5 dpc (A) and 18.5 dpc (B) embryos showing no apparent changes in stage specific cartilage and bone staining between control and Elavl1 cKO mice at these embryonic stages. Fore limb (C), Hind limb (D) from the 18.5 dpc stage embryos showing skeletal elements and maturation.

In addition, we measured bodyweight, lengths of tibia and femur at the indicated ages between control and Elavl1 cKO mice. There was no difference in the body weights between littermate controls and Elavl1 cKO animals (Fig-4A). In 8 weeks old male and female mice, the lengths of tibia and femur from hind limbs also did not differ significantly between controls and Elavl1 cKO mice (Fig-4B).

**Figure 4.**
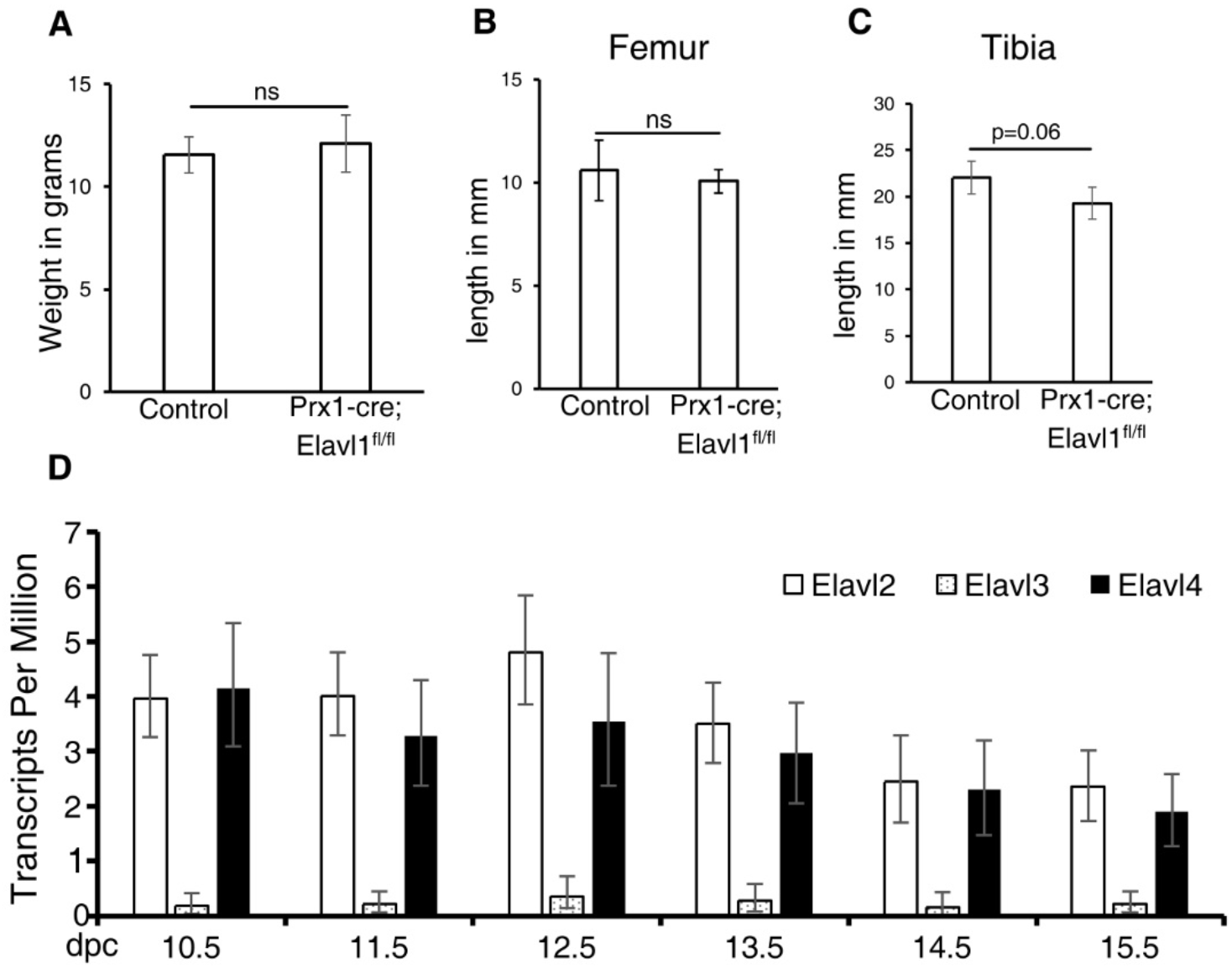
Body weight of 4-week-old male mice from control (N=3) and Elavl1cKO (N=3) groups. p= 0.6, ns: not significant. Femur (B) and tibia (C) lengths in the proximal-distal axis in 8-week-old male mice between control and Prx1-cre; Ealvl1^fl/fl^ mice, p= 0.59 (Femur) and p= 0.06 (tibia) ns: not significant. D. RNA-seq analysis of messenger RNA levels of Elavl2, Elavl3 and Elavl4 in limb RNA across developmental stages 10.5dpc-15.5 dpc indicating expression seen across these developmental stages.

Together, our results showed that Elavl1 is dispensable during appendicular skeletal development. Unlike other stem cell compartments including in the bone-marrow, intestine, and thymus wherein Elavl1 plays crucial roles during tissue development and homeostasis, deletion in limb progenitor cells did not result in disruption of skeletal patterning or morphological development. Deletion of Elavl1 globally during early development led to embryonic and extra embryonic developmental defects with embryonic lethality post E14.5 dpc. Targeted embryonic deletion of Elavl1 in epiblast cells using Sox2-cre also revealed significant changes during embryonic development affecting multiple tissues including spleen, skeletal tissues, lungs etc with no viable pups after birth^15^. However, tissues such as stomach and pancreas appeared normal in the absence of Elavl1 in conditional knockouts. Following tamoxifen inducible Rosa26Cre-ERT2 mediated deletion of Elavl1 in 8 weeks old mice, atrophy of multiple tissues was observed leading up to death within 10 days after administration of tamoxifen^16^. These studies indicated differential tissue susceptibility to Elavl1 depletion during development. In this study, Elavl1 gene was deleted conditionally and, in a tissue specific manner in the developing limb buds using Prx1 enhancer mediated cre expression for a more specific and localized deletion in developing limb buds and cranial mesoderm where high expression of Elavl1 was detected. Viable Elavl1 conditional knockouts were obtained in expected mendelian ratios during pre and postnatal development indicating the dispensability of Elavl1 during limb development. Further, developmental and molecular analyses of embryos also did not reveal any morphological or structural changes in the limb during or post development.

Elavl1 is a member of Elavl protein family which includes Elavl2, Elavl3 and Elavl4 (also known as HuR, HuB, HuC and HuD respectively). Unlike Elavl1 which is ubiquitously expressed, other Elavl family members showed high expression in nervous system with very little known about their developmental expression patterns and roles in other tissues^8,9^. Indeed, except Elavl3, all the other Elavl family members are expressed in developing limbs (Fig-4C). It is possible that absence of Elavl1 in the limb buds might be compensated due to redundant functions of other members of Elavl family members as Elavl2 and Elavl4 that are expressed in developing limbs during same embryonic stages. Elavl2 knockout leads to incomplete lethality and growth retardation in mice postnatally^23^ further showing neuronal independent functions for other Elavl1 family members. Further studies with compound mutations of Elavl family members will shed more light on the functions of specific members of this RNA binding protein family during development of various tissues.

## Acknowldgements

The research work reported in this paper was aided by grants K01AR069197 (to S.K.K from NIH/NIAMS), and R01GM135377 (to S.K.K from NIH/NIGMS).

## Notes

### Competing Interest Statement

The authors have declared no competing interest.

